# Tree-weighting for multi-study ensemble learners

**DOI:** 10.1101/698779

**Authors:** Maya Ramchandran, Prasad Patil, Giovanni Parmigiani

**Affiliations:** Department of Biostatistics, Harvard T.H. Chan School of Public Health; Department of Biostatistics and Computational Biology, Dana-Farber Cancer Institute

**Keywords:** Ensemble learning, Random Forests, Replicability, Multiple studies

## Abstract

Multi-study learning uses multiple training studies, separately trains classifiers on individual studies, and then forms ensembles with weights rewarding members with better cross-study prediction ability. This article considers novel weighting approaches for constructing tree-based ensemble learners in this setting. Using Random Forests as a single-study learner, we perform a comparison of either weighting each forest to form the ensemble, or extracting the individual trees trained by each Random Forest and weighting them directly. We consider weighting approaches that reward cross-study replicability within the training set. We find that incorporating multiple layers of ensembling in the training process increases the robustness of the resulting predictor. Furthermore, we explore the mechanisms by which the ensembling weights correspond to the internal structure of trees to shed light on the important features in determining the relationship between the Random Forests algorithm and the true outcome model. Finally, we apply our approach to genomic datasets and show that our method improves upon the basic multi-study learning paradigm.

## Introduction

With the increasing availability of multiple datasets that measure the same outcome and many of the same features, it is important to consider the totality of this information to train replicable predictors. Training and validating a predictor on a single study can often produce inflated reports of accuracy, especially when compared with results obtained when training and validating on different studies (1). The issue of replicability motivates the use of ensemble learning methods to combine the predictive power of multiple studies.

A cross-study learner (2) is an ensemble of learners trained separately on different studies, or Single Study Learners (SSL’s). An SSL can refer to any algorithm that produces a prediction model using a single study. The SSL’s are then combined using a specified weighting strategy, creating a single predictor that can be applied to external studies. Given *K* training datasets, we first train an SSL on each dataset. Denoting *Ŷ*_*k*_(*x*) as the vector of predictions that SSL_*k*_ makes on a new point *x*, and denoting *w*_*k*_, for *k* = 1, …, *K*, the weight given to SSL_*k*_ in the ensemble, the resulting prediction made by the multi-study ensemble can be expressed as:

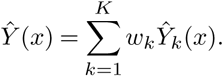

We investigate the utility of weighting schemes that enhance replicability and are robust to different population structures. This approach circumvents explicitly modeling the difference in feature-response relationships across studies. Using appropriately weighted ensembles of predictors each trained on a single study inherently takes into account study-specific features and creates a more replicable overall predictor that is agnostic to between-study heterogeneity in the marginal distributions of features and the conditional distribution of the outcome. Ensembles may be created using multiple SSL’s (for example, a neural net and linear regression), or a single type of SSL’s, which will be the setting explored in this paper.

The role of multiple studies in optimal ensembling strategies has been previously explored by Patil and Parmigiani (2). As a baseline, they considered the performance of a single learner trained by merging all the studies. They found that in the presence of inter-study heterogeneity, merging and meta-analysis performed worse than any of the cross-study methods considered. They additionally discovered that when considering homogeneous ensembles with Random Forests (3) as a SSL, ensembling always produced a better predictor than training one learner on the merged dataset, even in simulations where no between-study heterogeneity in the feature-outcome relationship was present. This is somewhat counter-intuitive, as the absence of between-study heterogeneity corresponds to a uniform true outcome model across studies (essentially, a single data-generating mechanism), yielding an equivalent optimal prediction rule for every dataset. It would appear in this case that merging all studies, thereby including more data points with the same feature-outcome relationship in training, would be most likely to result in a prediction rule that closely reflects the truth. Patil and Parmigiani found this to be the case for all considered learners other than Random Forests. For Random Forests, training SSL’s on each dataset and then ensembling produced better results than training one forest on the merged dataset, even though each dataset provides fewer samples for a single forest to learn the same prediction rule. This unexpected finding motivates our present exploration into the properties of Random Forest and optimal tree-weighting strategies.

The Random Forests algorithm (3) uses bagging (4) to train an ensemble of decision trees; two levels of randomization serve to make the trees less correlated and reduce the variance of the overall predictor. In the original algorithm, every tree within the Forest is given an equal weight in determining a prediction. We consider Random Forests as our SSL of interest to explore the utility of ensembling a group of SSL’s that are themselves already ensemble learners. We are particularly interested in the effect of extracting the trees trained by each SSL forest and weighting them directly, instead of simply assigning weights to each forest and giving all internal trees equal weighting. This corresponds to changing what is viewed as an SSL in the combination strategy; instead of considering forests, we now view the trees trained within each forest as our new SSL’s of interest. We consider weighting approaches that reward cross-study replicability within the training set. The motivation behind this approach is to allow for more control over the weights given to each tree in each Random Forest SSL in the global ensemble. Rewarding cross-study replicability on the tree level should increase the robustness of the overall ensemble to differing population parameters, such as between-study heterogeneity in the relationships between the features and the outcome and interactions between features in determining the outcome.

In this paper, we examine whether directly weighting trees improves on weighting forests, using simulations based on datasets including gene expression measurements and survival outcomes in ovarian cancer patients. We additionally apply our tree-based ensembling methods on other genomic multi-study and large-scale datasets.

## Methods

### Ovarian Cancer Datasets

CuratedOvarianData (5) from Bioconductor in R provides data for gene expression meta-analysis of patients with ovarian cancer. In this study, we use all 15 studies in CuratedOvarianData that provide survival information without any missing data in the features. The sample sizes of each dataset range from 42 to 510 subjects, and the fifteen datasets have 2,909 gene features in common. The gene expression features in each dataset are normalized for the analysis.

### Simulating datasets

To produce realistic simulations, we used feature vectors resampled from real data taken from CuratedOvarianData, to ensure that the predictors maintain realistic distributions and covariances. For N = 100 iterations for each set of generative parameters, we randomly separated the 15 datasets from CuratedOvarianData into *K* = 10 training and *V* = 5 validation sets. Per iteration, we reduced each dataset to the same randomly sampled 100 genes out of the 2909 comprising the intersection. 10 out of these 100 genes were then randomly chosen to create a linear data-generating model. Coefficients were chosen for each gene uniformly from [−5, −0.5] ∪ [0.5, 5], which ensured that each chosen gene has a non-zero contribution to the outcome. User-specified between-study heterogeneity in the feature-outcome relationship was then introduced through coefficient perturbation. These perturbation windows were chosen to explore both small- and large-scale between-study heterogeneity in line with the differences in feature-outcome relationships we observe in the actual data. Denoting **c** as the vector of uniformly generated coefficients and *l* as the level of heterogeneity to be introduced, the new coefficients were generated uniformly from [**c** −*l*, **c** + *l*]. In every simulation, 5 of the training datasets were given low levels of coefficient perturbation, the other 5 higher levels, and the validation sets an intermediate level. At baseline, these levels were set as .25, 1, and .4 respectively. The baseline values were chosen to introduce relatively small perturbations throughout, with the validation level representing a middle ground between the two levels used in training, in order to determine the effect of training on studies with both higher and lower levels of heterogeneity. The heterogeneity levels were varied through the course of the analysis to evaluate the performance of the ensembles; in all figures, the heterogeneity level corresponds to to the highest level assigned to 5 of the training sets. The low level for the other 5 training sets was held constant at .25, and the intermediate level given to the validation sets was half of the highest level. We used a location shift instead of a scale change so that the features could be affected differently by between-dataset variation. We then generated the outcome vector *Y*_*k*_ for each study *k* = 1, …15 conditional on the chosen subset of the observed predictors and the simulated coefficients.

We also included the option to add interaction terms between some of the features when generating the outcome. In these simulations, we considered two different scenarios regarding interactions: (1) Two datasets in the training set have interaction terms between features; two in the validation set contain interaction terms, and (2) Six datasets in the training set and two in the validation set have interaction terms. For all scenarios involving interaction terms, baseline heterogeneity parameters were fixed at moderately low levels.

We also employ modifications to this method when generating baseline simulations, as described in the Results section.

### Stacked regression weights

The purpose of this analysis is to develop ensembles of Random Forests SSL’s that improve prediction ability. To construct the weights given to each SSL in the ensemble, we used the multi-study version of the stacked regression method (2), which rewards cross-study generalizability of SSL’s in determining the ensembling weights. Stacked regression (6) provides a method for forming linear combinations of multiple predictors that improves on the performance of any single one. The weights given to each predictor in the combination are determined by cross-validation and least squares regression. Breiman’s paper (6) heuristically analyzes several modifications to least squares and finds that imposing a non-negativity constraint to the coefficients gives the best results. We considered variations to least squares regression, including Lasso and adding or eliminating an intercept, and found overall that using stacked regression with a Ridge constraint and intercept term produced the most robust ensemble on our setting. The Ridge constraint shrinks coefficients rather than directly zeroing some out; this appears to allow for greater generalizability of the resulting predictor. Setting the contributions of SSL’s directly to zero within training due to poor cross-validation performance may promote overfitting to the training datasets, as such SSL’s could provide value when faced with an observation arising from a new dataset with a potentially different structural features.

Our application of the stacked algorithm proceeds as follows: if we let *K* equal the number of training datasets available, we first train an SSL on each training set, yielding *K* different prediction functions. Denoting the predictions of SSL_*k*_ on Dataset_*k*_ for *k* = 1, …, *K* as *Ŷ*_*k*_, we stack the predictions of each SSL on every training dataset into one matrix, which here we will denote as **T**. 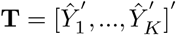, where *Ŷ*_*k*_ = [*Ŷ*_1*k*_ …*Ŷ*_*Kk*_]′ for *k* = 1, *…*, *K*; *Ŷ*_*ik*_ is the vector of predictions of SSL_*k*_ on dataset *i*, and *Ŷ*_*k*_ is the stacked vector of all predictions made by SSL_*k*_ on every training dataset. Denoting the total number of observations in all datasets as *N*, the dimension of T is *N* × *K*. We stack the true outcomes across all training studies into one vector, denoted as *Y*, whose dimension is *N* × 1. We then regress the observed outcomes stacked across all studies against the set of predictions, with a ridge constraint. The weights **w**_*stack*_ given to each SSL in the ensemble and are determined by solving:

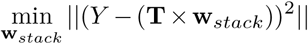

such that **w**_*stack*_ > 0 and 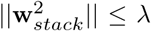, where *λ* is a hyperparameter that we optimize using the standard cross-validation procedure conducted by the `glmnet` package in R (7).

Throughout this paper, *Weighting Trees* will indicate individually weighting each tree using stacked regression with Ridge regularization and a nonnegativity constraint, while *Weighting Forests* will indicate using the same regularized stacking method on the predictions made by whole forests. The term *Unweighted* will refer to the ensemble created by giving every single-study forest equal weighting. The term *Merged* will refer to the predictor trained using a single learner on the single dataset formed from merging the 15 studies together. The difference between the Weighting Trees and Weighting Forests approaches is in what constitutes an SSL: individually weighting trees corresponds to first training a Random Forest on each study, then extracting the trees and treating each of the extracted trees as an SSL. If *m* is the number of trees per forest, the length of **w**_*stack*_ for the Weighting Trees approach is *K* × *m*, while for the Weighting Forests approach it is *K*. In some analyses we compare tree-level weights for Weighting Trees to the tree-level weights implied by Weighting Forests, which we define b y d ividing e ach forest-level weight from **w**_*stack*_ by *m*, as each tree within the *k*^*th*^ forest is implicitly assigned a weight of 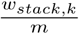.

### Number of trees per forest

In the implementation of Random Forest contained within the `randomForest` package in R (8), the default number of trees is set to 500 per forest. In general, the performance of Random Forest increases as a function of the number of trees before reaching a plateau; for this reason, the number of trees/forest is not typically considered to be a tuning parameter optimized through cross-validation. We found in our preliminary simulations that the relationship between approaches was conserved as the number of trees per ensemble increased; this is illustrated in Figure S2 from the supplement. In fact, we saw larger gains in predictive performance for the ensembling methods we evaluate when using greater numbers of trees per forest. For speed of computation, we used 10 trees per forest in all ensembles of our main analyses, but we anticipate that the improvements we observe are conservative representations of what is possible with larger forests. We trained the Merged learner with the same number of trees contained in the total weighted ensembles, in order to keep the total number of trees for each method equal. Typically, this means that there are 100 trees per ensemble, unless otherwise specified.

## Results

For all simulated results, averages are taken over 100 simulations at the specified parameters, with 95% confidence bands computed using the mean ±1.96× standard error.

### Baseline analyses

Figure 1A displays the percent variation in the outcome explained by interaction terms in the data generating mechanism as a function of increasing interaction strength. In this context, interaction strength represents the magnitude of the coefficients given to interaction terms in the linear model used to generate the outcome. The level of between-study heterogeneity does not affect the percent variation in the outcome, and there is a roughly linear relationship between interaction strength and percent variation across all heterogeneity levels considered.

**Fig. 1.**
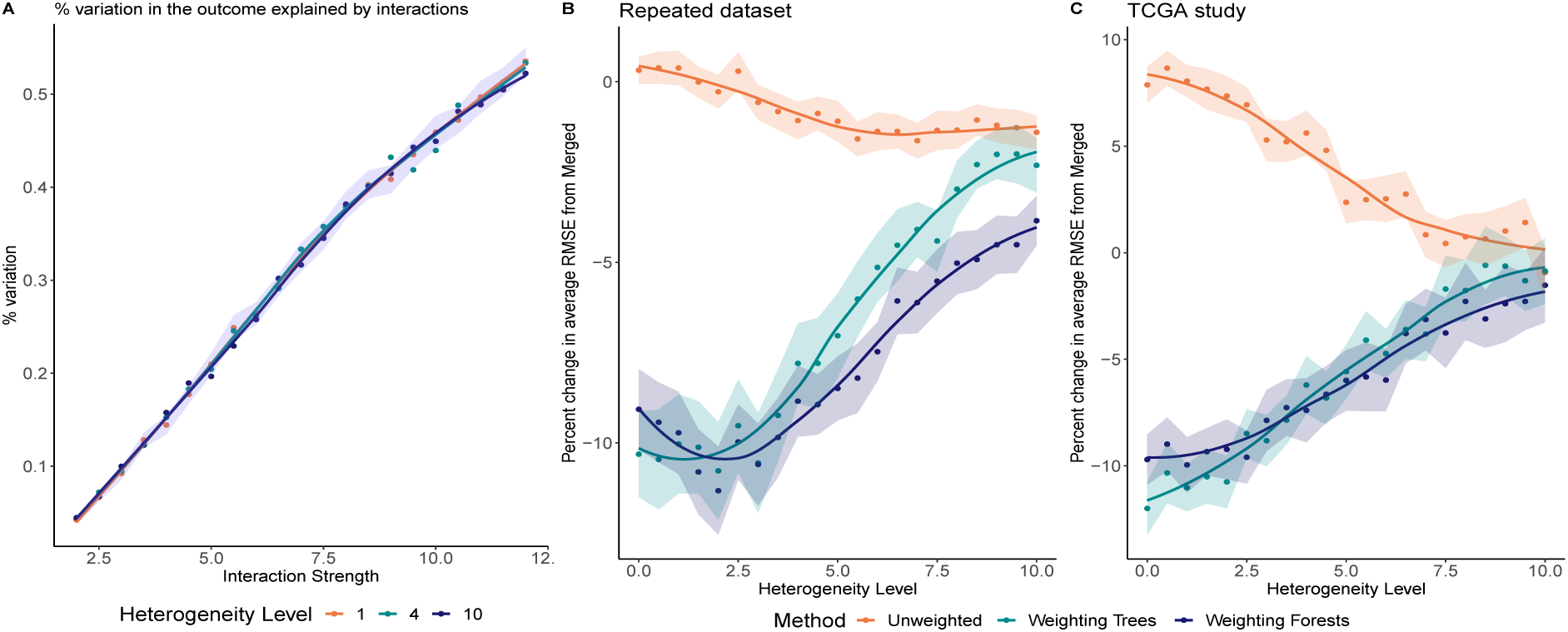
Baseline analyses of interaction strength and heterogeneity. **(A)** Percent variation in the outcome explained by interactions between features, as a function of interaction strength. **(B)**-**(C)** Percent change in average RMSE of each of the ensembling approaches compared to the Merged learner, as a function of between-study heterogeneity level. In **(B)** all training sets have identical feature distributions while in **(C)** the TCGA study is randomly split into 5 sub-datasets at every iteration for training and testing. Weighting Trees and Weighting Forests significantly improve upon Merged, with the difference in performance decreasing as heterogeneity i ncreases. Smoothing is applied to reduce simulation noise.

To preface analysis of panels B and C of Figure 1, we must first distinguish between “feature distribution heterogeneity” and “feature effect heterogeneity” (see also (9)). The former refers to changes in the distribution of features between datasets. The latter refers to changes in the relationship between the features and the outcome: in our simulations, it is captured by changes in the coefficients of the linear model used to generate the outcome, and is used throughout as the horizontal scale in the figures. At baseline, we explored removing feature distribution heterogeneity between datasets and evaluating the performance of the ensembles at differing levels of feature effect heterogeneity. Both types of heterogeneity may influence the utility of ensembling, so these simulations helped us more sharply analyze feature effect heterogeneity alone.

In the simulation used to generate Figure 1B, we set *K* = 10. Each of the *K* studies has the same sample size and the same feature set obtained by repeating a single randomly chosen ovarian cancer dataset *K* times, thus removing feature distribution heterogeneity. Each iteration has a different level of feature effect heterogeneity. In each we train the Merged learner and the SSL’s and evaluate them on 5 independent datasets not used in training. The Weighting Trees and Weighting Forests approaches perform better than the Merged learner across all levels of heterogeneity considered, while the Unweighted method generally only slightly improves on the Merged. Training an SSL on each set of observations and upweighting those learners with better cross-study validation performance (which at level 0 is akin to cross-validation) produces markedly better results. The difference is most pronounced at levels of feature effect heterogeneity around 2, where it exceeds 10%, and is substantial at level 0. As the level of heterogeneity rises, the underlying RMSE increases, and the difference in performance among approaches decreases. We also evaluated the performance of a single learner trained on one copy of the dataset chosen to be repeated in the training set, and found that in the presence of no feature distribution heterogeneity, the ensembles using the stacking weights trained on the repeated training set outperform this single learner.

The dataset used to train the Merged includes each predictor profile in the training dataset 10 times, a somewhat artificial setting. For Figure 1C, in each iteration, we split the TCGA study (comprising 510 subjects) into 5 equally sized pseudo-datasets. We used 4 to train the merged learner and SSL’s, and kept the 5th to test the resulting ensembles. This simulation helped us investigate whether differences in the the distribution between smaller sub-datasets and the whole dataset would be sufficient to result in a better performance of ensembling approaches compared to merging. Both the Weighting Trees and Weighting Forests approaches outperform the Merged at all heterogeneity levels, while the Unweighted ensemble results in a decrease in performance.

### Overall performance of ensembling approaches

Our next simulations include both feature distribution and feature effect heterogeneity. Figure 2 displays average RMSE’s over 100 iterations for the various ensembling approaches for three different scenarios. Out of 100 total features present, 10 features affect the outcome; the outcome is continuous. Panel

**Fig. 2.**
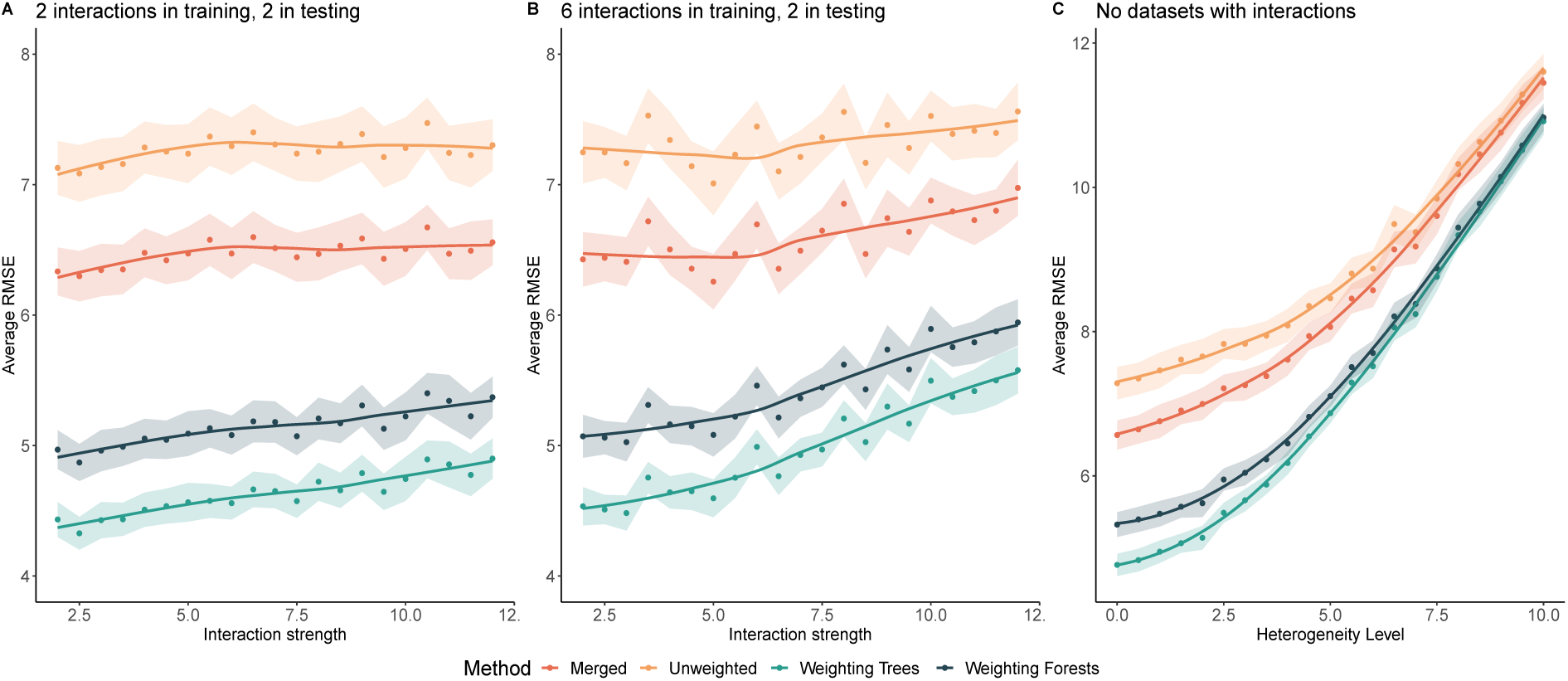
Average RMSE’s of ensembling approaches (color labeled) across different data-generating scenarios, as a function of increasing interaction strength or heterogeneity. **(A)** 2 datasets with interaction terms between features in the outcome-generating generating mechanism are included in the training set, and 2 are included in the testing set. **(B)** 6 datasets with interactions are included in the training set, 2 in the testing set. **(C)** No datasets with interaction terms are included in either training or testing, and performance is evaluated for increasing feature effect heterogeneity. For all three scenarios, and across parameter levels, Weighting Trees outperforms all other approaches.

A corresponds to interaction scenario (1) as described in the Simulating Datasets section, while panel B corresponds to scenario (2). Panel C considers no interactions and considers the effect of increasing the level of feature effect heterogeneity.

Overall, both Weighting Trees and Weigthing Forests perform significantly better than either of the simpler approaches considered as baseline comparisons. Interestingly, ensembling forests is only better than merging when we suitably weigh each forest or tree, which we see because the Unweighted approach consistently performs worse than the Merged. The Merged and Unweighted each give the same overall weight to each tree (1/100), but they differ in the number of candidate data points available to form a bootstrap sample to train each tree. We can extrapolate that training each tree on the merged dataset produces a predictor with better generalizability than restricting each tree to a single dataset and using simple averaging to combine the resulting forests. This motivates the necessity of using weighting approaches that reward cross-study replicability when constructing an ensemble of ensembles. Otherwise, it may be both more efficient and accurate to rely on the ensembling produced within a single Random Forest.

Furthermore, the Weighting Trees strategy consistently performs better than Weighting Forests as well as the other baseline approaches. This follows the intuition that we can create a more robust ensemble by rewarding cross-study replicability at the individual tree level, by considering the strengths and weaknesses of individual trees rather than forests in terms of out-of-sample cross-study validation performance within the studies in the training collection.

A comparison of panels A and B from Figure 2 indicates that including more datasets in which interactions are present in training does not generally improve the performance of any of the ensembling constructions considered; additionally, there is more variability around the average trend line. This suggests there is a balance in creating an optimally heterogeneous set of training studies for ensembling, and that the presence of heterogeneity could be more important than the sample size.

Figure 2C demonstrates that as the level of feature effect heterogeneity increases, the differences in performance of the ensembling approaches we considered decreases until they become virtually indistinguishable. At the lowest level of heterogeneity, we observe a significant separation between Weighting Trees, Weighting Forests and the simpler approaches. This suggests that in our data, there is intrinsic cross-study heterogeneity in the distribution of the features, that motivates utilizing replicability weights over merging or simple averaging, supporting the conclusions in the baseline simulations.

### Tree structure and variable importance measures

One of our primary interests in this study was to explore how the trees trained by Random Forest capture the relationship between the variables and the outcome. We specifically wanted to elucidate which features, or variables, are important to weighting schemes that reward replicability.

Table 1 displays results about how tree structure and variable importance measures are related to the weights given to each tree in each ensembling approach. The ‘true’ variables are the variables chosen to linearly combine to generate the outcome in each simulation. All averages are taken over 100 iterations of the simulation; 2 datasets in both the training and testing group contain interaction terms. For all ensembles, there are 9 total variables per tree; in the data generating mechanism, 3 variables out of the 10 associated with the outcome are involved in interaction terms.

**Table 1.** Tree structure summaries and variable importance measures for each ensembling approach, by tree weight. The left three columns are average tree-level measures. The rightmost column measures the proportion of the total sum of variable importance scores attributed to true variables. Variable importance is computed as follows: for each tree, the prediction error on the out-of-bag portion of the data is recorded (using MSE for regression). Then the same is done after permuting each predictor variable. The difference between the two are then averaged over all trees, and normalized by the standard deviation of the differences.

| Tree weight percentile | Average tree weight | Freq. of true variables/tree | # of variables in interactions | Proportion of total varImp by true vars. |
| --- | --- | --- | --- | --- |
| Weighting Trees |  |  |  |  |
| 90-100% | **0.068**<br>(0.0675, 0.0685) | **0.402**<br>(0.400, 0.404) | **1.459**<br>(1.443, 1.475) | **0.699**<br>(0.671, 0.727) |
| 0-10% | **0.001**<br>(0.001, 0.001) | **0.143**<br>(0.141, 0.145) | **0.48**<br>(0.468, 0.492) | **0.502**<br>(0.477, 0.527) |
| 0-100% | **0.026**<br>(0.0259, 0.0261) | **0.265**<br>(0.264, 0.266) | **0.948**<br>(0.937, 0.959) | **0.655**<br>(0.627, 0.683) |
| Weighting Forests |  |  |  |  |
| 90-100% | **0.0463**<br>(0.0458, 0.0468) | **0.368**<br>(0.364, 0.372) | **1.348**<br>(1.326, 1.37) | **0.704**<br>(0.675, 0.733) |
| 0-10% | **0.0091**<br>(0.0089, 0.0093) | **0.171**<br>(0.169, 0.173) | **0.571**<br>(0.558, 0.584) | **0.512**<br>(0.486, 0.538) |
| 0-100% | **0.0256**<br>(0.0254, 0.0258) | **0.265**<br>(0.264, 0.266) | **0.948**<br>(0.937, 0.959) | **0.655**<br>(0.627, 0.683) |
| Merged |  |  |  |  |
| 0-100% | **0.01** | **0.504**<br>(0.502, 0.506) | **1.889**<br>(1.869, 1.909) | **0.624**<br>(0.600, 0.648) |

For both tree-based and forest-based ensembles, trees with larger weights also have a higher percentage of true variables, number of variables involved in interactions, and true variables with higher variable importance. Interestingly, the frequency of true variables and true interaction variables in trees in the Merged learner is higher than in the top decile for either Weighting Forests or Weighting Trees. Yet the Merged has worse overall performance than both ensembling approaches. We speculate that these metrics of tree-level variable importance are imperfect markers of cross-study generalizability, as they do not address the stability of a variable’s effect which is required for replicability. The true variables in each iteration of the simulation are shared across all studies, so the frequency at which such variables appear within trees is not intrinsically a study-specific feature. Similarly, the variables chosen to be involved in interaction terms are shared across all studies chosen to contain interactions, trees trained on such studies share similarities across studies. However, the coefficients do vary, and cross-study learners attempt to build predictors that are more robust to this variation.

The top decile of tree-based ensembles and forest-based ensembles contain on average nearly half of the 3 variables involved in interactions, with the tree-based average slightly higher than that for the forest-based ensemble. This is significantly higher than either the total average or the average within the lowest decile for either method, indicating that when the true relationship between the features involves interactions between variables, trees containing such variables have markedly increased cross-study generalizability and are therefore given higher weights within the ensemble.

### Distribution of tree-level weights

In general, the distributions of the weights given to individual trees have a similar location, but are more dispersed than those given to forests, and have a more pronounced upper tail. From Table 1, we see that for the Weighting Trees approach, the lowest decile of trees have weights that are 10-fold lower compared to the merged, while the top decile is approximately 7 times higher. The Weighting Forests method does not down-weigh the lowest decile or up-weight the top decile to the same extent. To produce optimal results, there is a balance between giving all trees equal weight and having too big of a weight discrepancy between trees. Overall, the relationship between tree weight and variable importance features is very similar for tree-based ensembles and forest-based ensembles. Forest-based ensembles don’t allow for control over tree-level weights, resulting in a narrower overall distribution. The benefits afforded by a few trees are potentially averaged out in the performance of the corresponding forest and potentially not rewarded optimally.

Figure 3 illustrates this. The overall distribution of weights given to trees within forest-based ensembles is narrower than that for tree-based ensembles, although both have relatively similar medians and means. The dotted red line at *y* = 0.01 shows the weight that would be given to each tree if all were equally weighted, as in the Merged and Unweighted. Both tree- and forest-weighting approaches produce distributions centered above *y* = 0.01, as we implement stacking without a normalization of weights. The distribution of the difference between the weight given to the same tree by the Weighting Forests and Weighting Trees approaches is centered around zero, but displays a pronounced lower tail, suggesting that Weighting Trees may identify a relatively small subset of trees for up-weighting, while reducing the weight of most others, compared to Weighting Forests. Consistently, as displayed in Table 1, the top decile of the Weighting Forests distribution is on average smaller than that for Weighting Trees while the overall means are roughly equal. Essentially, trees that are deemed by the Weighted Trees approach to be among the least beneficial in the ensemble may still receive moderate or high weight if the whole forest is weighted together. This likely leads to the decrease in performance we see through-out.

**Fig. 3.**
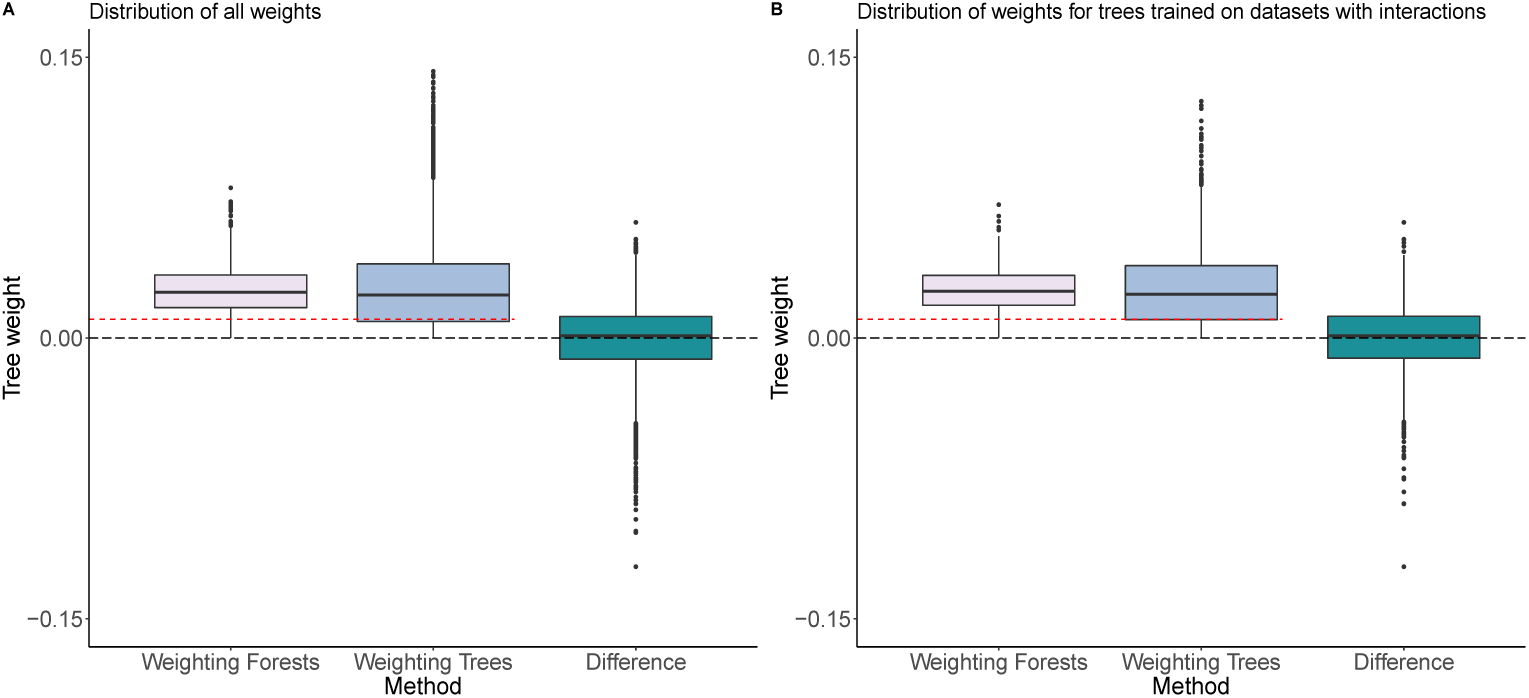
Distribution of tree-level weights given to ensembles trained using the Weighting Trees or Weighting Forests methods, as well as their difference, defined as the weight given by Weigting Forests minus that given by Weighting Trees; 2 of the training sets contain interaction terms in the outcome-generating mechanism. Tree-level weights for Weighting Forests are obtained by dividing the forest-level weight returned by the stacking algorithm by the number of trees per forest. Correspondingly, each point in Weighting Forests represents the value of the weight given to 10 trees. **(A)** Distribution of the weights given to all trees in each ensemble. **(B)** Distribution of the weights given only to trees trained on datasets containing interaction terms. The dashed red line at *y* = .01 represents the weight given to every tree within the Merged and Unweighted.

Figure 3B displays the distribution of weights for only those trees within each ensemble that were trained on datasets containing interaction terms in the outcome-generating mechanism. The general distribution for these weights is similar to the distribution of all weights within the ensemble. This is an interesting result, with multiple potential causes. The Random Forest algorithm specifies that only a randomly chosen subset of all features are available at each split during the training of each tree. Only some of the trees within forests trained on datasets with interaction terms actually contain the features involved in interactions; Table 1 illustrates that such trees tend to be up-weighted, but do not comprise the entirety of their corresponding forests. Moreover, the ridge constraint in the stacking algorithm shrinks the coefficients towards zero, so no given tree can ever be drastically upweighted compared to the rest. Determinants such as these result in a similar overall weighting distribution as seen in the whole ensemble.

Furthermore, there are several factors (both measured and unmeasured) that contribute to improving cross-study generalizability of individual trees, not only the presence of variables involved in interaction terms. It appears that the interplay between all such components is paramount in determining the weight of each tree, and that isolating the presence or absence of any single one will not carry significant influence.

### The Combined approach

Our simulations so far focused specifically on the comparison between weighting trees and weighting forests, using learning from the merged data as a benchmark. Table 1 suggests that trees identified by single study learners capture different features compared to trees trained on merged data sets. Having established that weighting individual trees improves prediction ability, we can now ask whether this concept can be extended to collections of trees trained both on single studies and merged data. Table 1 reveals that the Merged approach can more frequently identify true predictive features. It is possible that useful trees may be identified often in the merged training, but may not be adequately leveraged because of equal weighting.

To explore this, we implemented a Combined Weighting Trees method (Combined for short), which computes stacked regression weights with a ridge constraint on the union of the 100 trees trained on the merged dataset and the 10 ×10 = 100 trees trained by the SSL approach; the resulting ensemble comprises 200 trees. We then reconsider the simulation setting of Figure1C.

Figure 4 summarizes the results. The Combined approach improves upon all of the other ensembles for most heterogeneity levels considered. The greatest gap is observed at 0, when no feature effect heterogeneity is included. This separation in performance decreases with heterogeneity increases. As heterogeneity increases, so does the importance of modeling both the changes in features distribution and relationship between the features and the outcome; this results in a decreasing difference between including trees trained on the Merged in the Combined approach and simply Weighting Trees or Weighting Forests. Subsequently, when tested in the multi-study simulation setting described previously, the Combined approach did not improve upon the weighting trees approach, possibly because of the more central role of feature distribution heterogeneity.

**Fig. 4.**
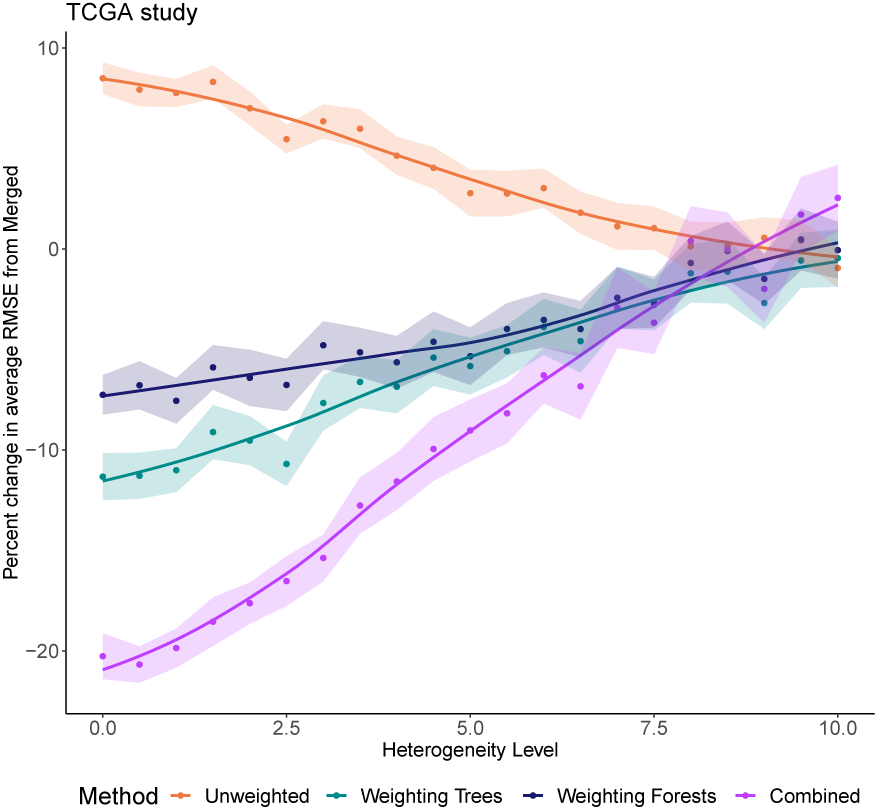
Performance of the Combined approach compared to the other ensembling methods on the TCGA study; the TCGA study is randomly split into 5 sub-datasets at every iteration for training and testing. The Combined significantly improves on the rest for lower levels of heterogeneity between the sub-datasets.

## Data Applications

To explore the performance of these classifiers on real data with natural feature-outcome relationships, we considered relatively high-dimensional multi-study datasets in which we had reason to believe there was inter-study heterogeneity and interactions between features. Gene expression and clinical data were natural candidates.

### Multistudy setting: Breast Cancer datasets

We first considered multi-study datasets; CuratedBreastData from Bioconductor in R provides gene expression and clinical data for 34 studies following patients with breast cancer (10). All of the relevant clinical information collected in the datasets is binary; there were five total studies that measured Overall Survival (OS) of patients, which we chose as our clinical outcome of interest. OS is defined as 1 if the patient survives to the end of the study period, and 0 otherwise. Study periods may vary, resulting in a further source of heterogeneity, affecting feature effects. The sample sizes range from 14 to 118, with a total of 336 subjects across studies. The five datasets have 76 gene features in common and 47 clinical features with fewer than 10 missing datapoints across studies. These 123 variables were used as features in the analysis. We tested the performance of the ensembling approaches when predicting OS while training on four studies and validating on the fifth, using log loss as our metric of success. Since `randomForest` trains a different set of trees at each iteration, we replicated the method 100 times to obtain average log loss values and standard errors.

The results are pictured in Figure 5A.1-A.2. The Weighting Trees approach is the overall best for predicting OS. The Merged learner performs several orders of magnitude worse than the rest, in both relative and absolute terms. Conversely, the Unweighted, Weighting Trees, and Weighting Forests methods have low absolute prediction Log Loss. This agrees with the results from simulations using binary outcomes shown in Figure S1 from the supplement.

**Fig. 5.**
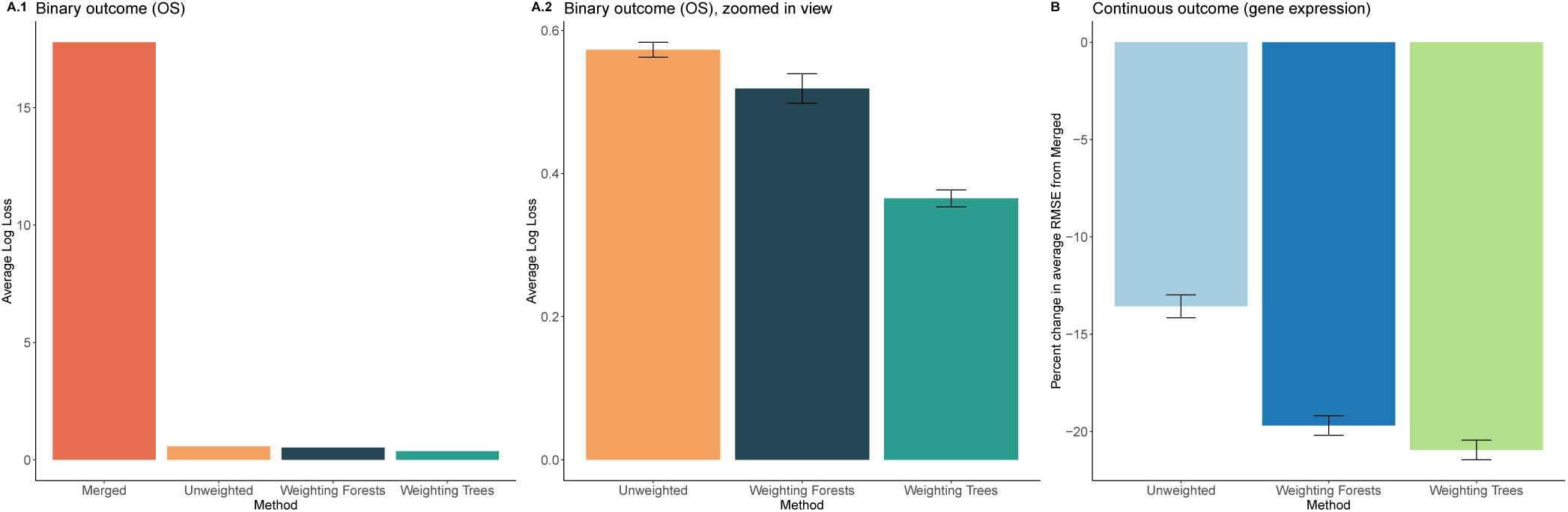
Performance of ensembling approaches on the breast cancer datasets in the multistudy setting, with associated 95% confidence intervals. **(A.1)** Average percent change in prediction Log Loss from the Merged for each of the ensembling approaches on the binary outcome variable, Overall Survival (OS). Confidence intervals were obtained by training each of the ensembling approaches 100 times, with differences in performance across iterations induced by the randomization within the Random Forest algorithm. **(A.2)** A view of panel A.1, without the Merged learner to improve scaling, so differences between the ensembles can be clearly visualized. **(B)** Average percent change in RMSE from the Merged when predicting expression levels for each of the top 500 variable genes given the rest of the gene expression data. The standard errors were therefore computed over 500 samples, as opposed to the 100 in panel A.2.

We also evaluated the performance of the ensembles on a continuous outcome. As there was no continuous clinical variable, we considered the task of predicting gene expression levels. Clinical decisions are often informed by expression levels of certain genes, thus predicting these levels when they may be missing can be useful in determining treatment. We used six studies in this analysis. The sample sizes range from 21 to 195 subjects, with 511 total across studies. The six datasets have 1,312 gene features in common. We predicted the expression level of one gene given the rest of the expression data, for the 500 most variable genes included in the features. Our performance metric was the percent change in average RMSE from the merged learner. The results, averaged across all 500 genes tested with associated 95% confidence intervals, are pictured in Figure 5B. The general relationships between the performance of the ensembling approaches follow what we see in the discrete outcome case, with the Weighting Trees approach outperforming the rest. It is slightly superior to Weighting Forests, and both surpass the Unweighted. All three ensembles significantly improve upon the Merged learner; this slightly differs from the results seen in the simulations, in which the Merged approach typically outperforms the Unweighted. This difference is likely due to different composition of the gene expression data in the breast cancer studies compared to that in the simulations, and provides more motivation for using ensembling over merging when dealing with multiple datasets.

### Single study setting: Breast Cancer datasets

We also considered the use of ensembling methods on large single datasets. The MetaGxBreast package from Bioconductor in R provides a collection of Breast Cancer Transcriptomic Datasets that are part of the MetaGxData package compendium (11). For this analysis, we used the GSE25066 study, which contains 508 subjects, and the Metabric study, which contains 1989 subjects; in both datasets, we included the 100 gene features that exhibit the most variability, as well as any clinical features with complete data. For the GSE25066 study, we looked at two outcomes: DMFS (Distant Metastasis Free Survival) status, and days to DMFS. Days to DMFS describes the number of days from diagnosis to appearance of a distant metastasis. A distant metastasis refers to cancer that has spread from the original (primary) tumor to distant organs or distant lymph nodes. DMFS status is a binary variable that describes whether or not appearance of distant meastasis occurs during the study period. For the Metabric study, we considered the outcome Days to Death, which measures days from diagnosis to death, if death occurs within the study period. Patients within the study with missing values (either due to dropout or because death did not occur during the study period) were omitted from the analysis, yielding 1989 patients with complete data.

When constructing ensembles using the GSE25066 study, we split the dataset into five randomly chosen, equally sized subdatasets, trained on four, and tested on the fifth. The results for the two outcomes considered for 100 iterations of such splits are shown in Figure 6A-B. For the Metabric study, we split the dataset into 11 pseudo-datasets, trained on 10, and tested on the 11*th*. The results for 100 iterations of such splits are shown in Figure 6C.

**Fig. 6.**
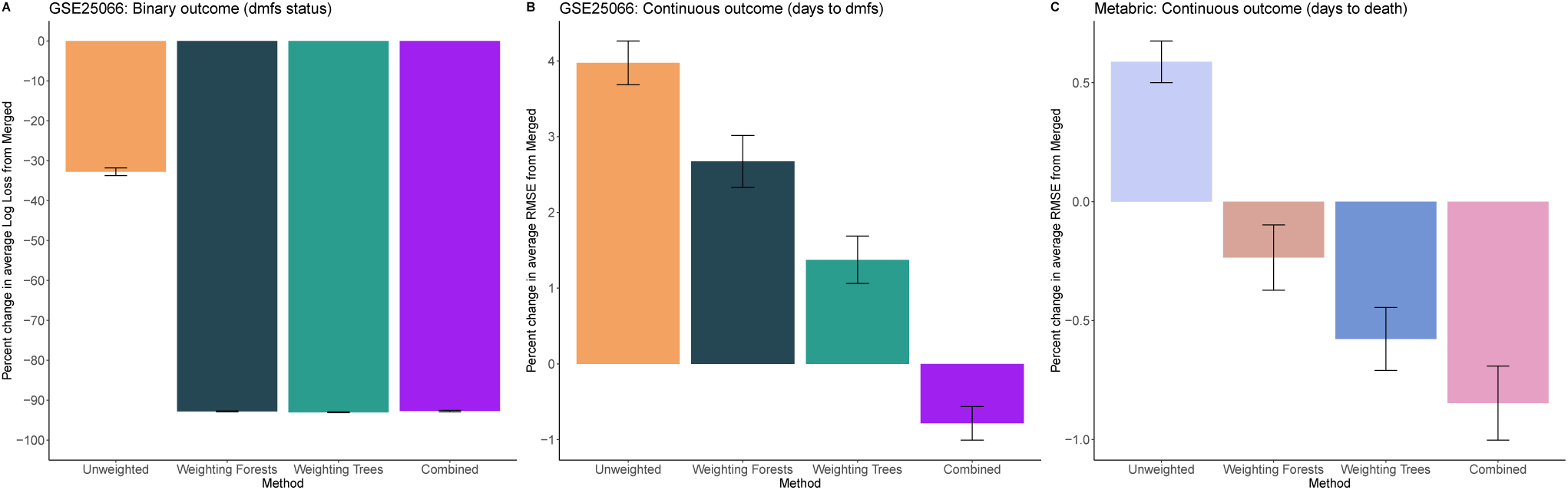
Performance of the different ensembling approaches on the breast cancer datasets in the single study setting, with associated 95% confidence intervals. Confidence intervals for all panels were obtained by iterating through the training and testing process 100 times. **(A)** Average percent change in prediction Log Loss from the Merged for each of the ensembling approaches on the binary outcome variable, Distant Metastasis Free Survival (DMFS) status. For every iteration, the GSE25066 Breast Cancer dataset (508 patients) was randomly split into 5 equally sized sub-datasets; 4 were then randomly chosen for training, and the 5^*th*^ was used for testing. **(B)** Average percent change in RMSE from the Merged when predicting the continuous outcome variable, days to DMFS, using the GSE25066 Breast Cancer dataset in the same manner as described in (A). **(C)** Average percent change in RMSE from the Merged when predicting the continuous outcome variable, days to death. For every iteration, the Metabric Breast Cancer dataset (1989 patients) was randomly split into 11 equally sized sub-datasets; 10 were then randomly chosen for training, and the 11^*th*^ was used for testing.

As evidenced by Figure 6A, all ensembling methods vastly outperform the Merged when predicting a binary outcome in the GSE25066 dataset; the three approaches that use cross-study weights are almost twice as accurate, while the Unweighted has more modest gains. The differences between the three top performers are negligible, indicating that for a binary dataset, using any of the weighting approaches that reward generalizability of trees or forests is sufficient to improve prediction ability by the same amount. The near equality of Weighting Trees and Weighting forests mirror the pattern seen in the simulations shown in Figure 1; however, the Combined does not improve upon either, unlike the simulations in Figure 4. This indicates that the SSL’s are able to capture enough of the whole dataset structure that the addition of trees trained within the Merged does not significantly improve performance.

Figure 6B shows the results from using the continuous outcome, Days to DMFS, in the GSE25066 dataset. Here, only the Combined approach described in the simulations section yields a better predictor than the Merged, and the improvement itself is only of the order of 1%. All other ensembles perform slightly worse than the Merged, with Weighting Trees producing the smallest difference. Containing 508 observations, the GSE25066 dataset is relatively large compared to many other gene expression and clinical datasets, which can typically be more on the order of 100-200 observations. However, we also wanted to determine whether the splitting method would be more appropriate on much larger datasets when the aim is to predict a continuous outcome.

To this end, we tested the performance of the ensembles on the Metabric dataset, which contains 1989 observations with non-missing outcome data (days to death); the results are displayed in Figure 6C. Here, all cross-study ensembling approaches slightly outperform the Merged, with the improvements all under 1%. Again, the Combined approach is the best of all methods considered, but unlike in the GSE25066 dataset, the Weighting Trees and Weighting Forests approaches yield marginally better predictions than the Merged. Although the range of improvement is not high, these results suggest that as the dataset gets larger, there is more benefit to using all cross-study ensembling. Overall, regardless of the size of the dataset or the type of outcome, the Combined ensemble always outperforms the Merged; this highlights its utility in the single dataset setting, particularly when dealing with a binary outcome.

The difference between the binary and continuous outcome results for the same dataset indicate the importance of the structure of the feature-outcome response in determining the performance of the ensembling approaches. We saw for the simulations that as the level of heterogeneity increased, all approaches converged; it appears that presence of considerable heterogeneity in the relationship between the features and outcome across the observations in the full dataset negatively impacts the performance of the ensembling approaches when the sub-datasets are chosen randomly. This applies to both datasets analyzed in this section.

## Discussion

In this paper we proposed and investigated a methodology for building ensembles of trees trained on multiple studies to achieve robustness to cross-study heterogeneity. Patil and Parmigiani (2) introduced a general multi-study learning architecture that uses stacking to combine predictors build on separate studies. In this setting, using Random Forests as a single-study learner, we compared weighting each forest to form the ensemble, as in (2), to extracting the individual trees trained by each Random Forest and weighting them directly. Our methodology extends the methods in (2).

Our results broadly indicate that Weighting Trees is often more effective than Weighting Forests, sometimes by a considerable margin. In turn, this suggests that more efficient families of cross-study learners can be constructed by “unpacking” learners that are themselves based on ensembles, and weighting the individual components directly.

Our results may depend on the specific implementation of the stacking algorithm used to reweight the trees. We show results using ridge regression stacking, but other approaches to regularization would be worth exploring.

Our strategy can be applied whether or not the training sample is naturally divisible into different studies or subpopulations. We investigated this case as a limiting case when heterogeneity is close to 0. The findings from our baseline analysis (see Figure 1) suggest that for larger datasets, it may be advantageous to train SSL’s on subsections and ensemble using stacking weights that reward prediction across different sections of the data, rather than simply training a learner on the whole dataset. Focusing SSL’s on fewer observations may allow the learners to capture more of the specific features of the dataset, which can then be combined in a way that promotes cross-study generalizability. Simply training one learner on the entire dataset may ignore potential heterogeneity in feature distributions between subsets of observations.

The results of the single study data example (see Figure 6) suggest that any weighting approach that rewards cross-study generalizability may be used in constructing ensembles for use on datasets with a binary outcome that are large enough to reasonably split into sub-datasets, as they vastly outperform a single learner trained on the entire dataset. Further work should be done in developing methods to determine advantageous groupings of observations based on sources of heterogeneity in the joint distribution of the predictive features. Since there are only two levels within a binary outcome, nuanced differences in the feature-outcome relationship across observations are harder to detect, as the amount of heterogeneity that maps to a particular label is significantly larger than that which would map to a particular continuous outcome value. This may be the reason for the substantial increase in performance of the weighted ensembles over the Merged even when the sub-datasets are randomly chosen.

For a continuous outcome, an important distinction to make between the simulations and data applications is that in the simulations, the outcome is defined to be linearly related to the features, while we don’t know whether this is true in the data applications. In the single-study setting, creating pseudo-studies generates simpler trees, which may be either advantageous or disadvantageous depending upon the complexity of the relationship between the covariates and outcome. In the simulations, in which this relationship is very clearly defined, generating simpler trees produces significant improvements in ensemble predictions. However, we see more ambiguous results in the continuous outcome single study data examples (6B-C); one possibility is that in this case, the covariate-outcome relationship is too complex to afford the advantages of simpler trees, resulting in only marginal gains in ensembling over merging.

Trees identified by single study learners capture different features compared to trees trained on merged data sets. We explored whether larger collections of trees including those trained both on single studies and merged data would provide improvements. This combined approach is advantageous when heterogeneity in the joint distribution of the features is low, particularly in the single study setting. Interestingly, some of these gains persist when there is moderate heterogeneity in the relationship between features and outcomes. This is intriguing and deserves further study, as the importance of modeling effect heterogeneity is paramount, and trees that do not take this into account are unlikely to promote generalizability to new datasets. Overall, in the single study setting, the addition of trees that do not model substudy heterogeneity and take into account the structure of the entire dataset enhances the performance of the trees within the SSL’s when no external between-study heterogeneity is added to the sub-studies. This suggests that the combined approach is a valuable candidate when considering ensemble methods for large single datasets.

## Reproducibility

Code and instructions to reproduce analyses are at https://github.com/m-ramchandran/tree-weighting.

## ACKNOWLEDGEMENTS

We thank Matt Ploenzke and Lorenzo Trippa for useful suggestions.

Maya Ramchandran and Prasad Patil were supported by NIH Cancer Training Grant (T32CA009337). Giovanni Parmigiani and Prasad Patil were supported by grant NSF-DMS 1810829. Giovanni Parmigiani was also supported by grant NIH-NCI 4P30CA006516-51.

