## Supplement for "Tree-weighting for multi-study ensemble learners"

July 10, 2019

[1] Department of Biostatistics, Harvard T.H. Chan School of Public Health
[2] Department of Biostatistics and Computational Biology, Dana-Farber Cancer Institute

This document serves as a supplement to the main article “Tree-weighting for multi-study ensemble learners”. Here, we present two supplementary simulations: the first explores the efficacy of ensembling approaches on a binary outcome, and the second evaluates the effect of increasing the number of trees per forest on performance using a continuous outcome.

### Binary outcome simulation

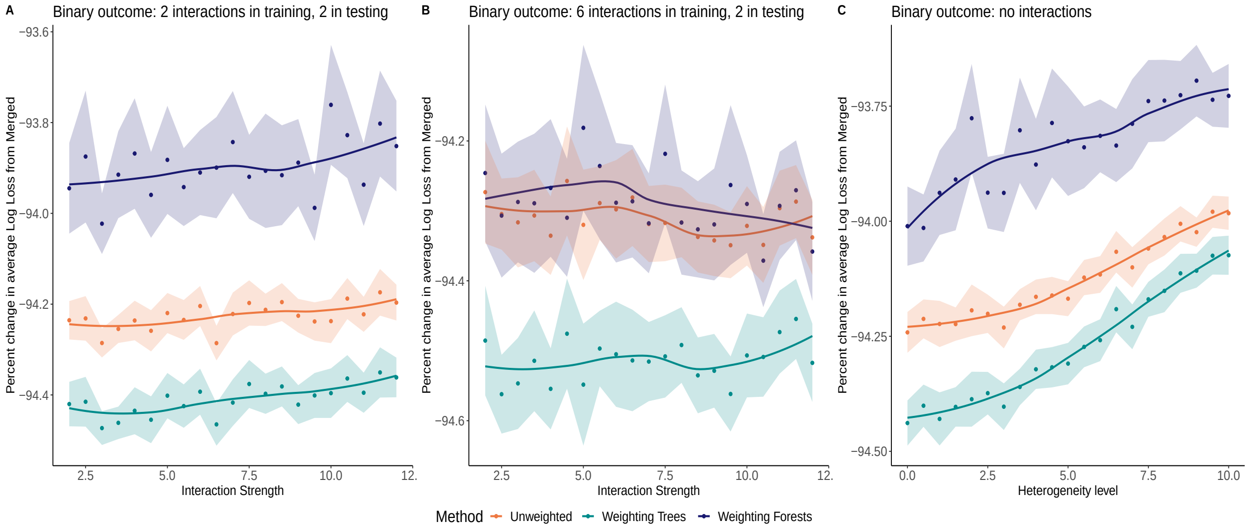


Figure S1: Average percent change in prediction Log Loss from the Merged for each of the ensembling approaches (color labeled) on a binary outcome variable, as a function of increasing interaction strength or heterogeneity. **(A)** 2 datasets with interaction terms between features in the outcome-generating generating mechanism are included in the training set, and 2 are included in the testing set. **(B)** 6 datasets with interactions are included in the training set, 2 in the testing set. **(C)** No datasets with interaction terms are included in either training or testing, and performance is evaluated for increasing feature effect heterogeneity. For all three scenarios, and across parameter levels, Weighting Trees outperforms all other approaches, and all three ensembling approaches vastly outperform the Merged.

Figure S1 displays average percent change in prediction Log Loss from the Merged over 100 iterations for the various ensembling approaches for three different scenarios, when the outcome is binary. Out of 100 total features present, 10 features affect the outcome; the outcome is continuous. Panel A corresponds to interaction scenario (1) as described in the Simulating Datasets section of the main text, while panel B corresponds to scenario (2). Panel C considers no interactions and considers the effect of increasing the level of feature effect heterogeneity. The datasets were generated in the same manner as described in the Simulating Datasets section, with one exception: the generated continuous outcome was dichotomized based on quantile, with outcomes values falling above the $75^{\text{th}}$ percentile being assigned a value of 1, and 0 otherwise. This created a binary outcome for the analysis.

In all three scenarios considered, all three ensembling approaches almost double their improvements over the Merged, with Weighting Trees outperforming the others throughout. In panels A and C, there are clear distinctions in the performances of each approach - interestingly, the Unweighted ensemble surpasses Weighting Forests, which is not a result we see in the continuous outcome setting. In panel B, the Weighting Forests and Unweighted approaches are nearly indistinguishable, while Weighting Forests exceeds them both.

### Increasing the number of trees in the ensemble

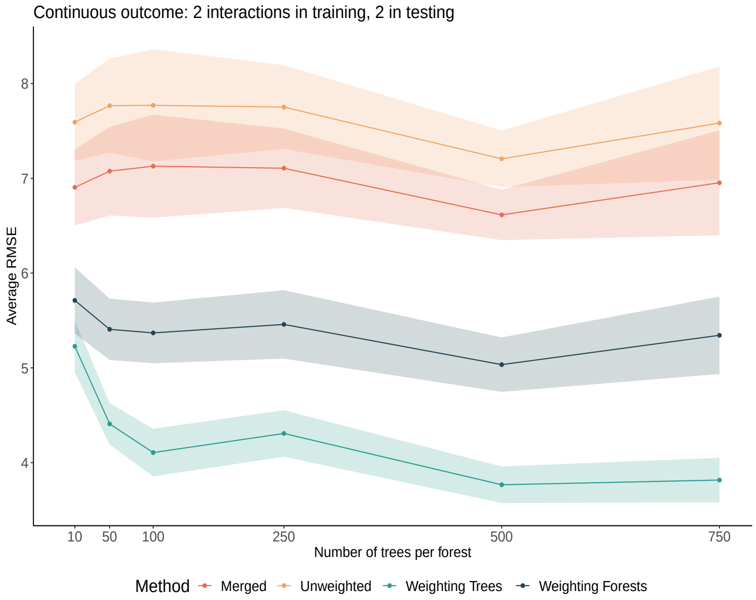


Figure S2: Average RMSE's of ensembling approaches (color labeled) on a continuous outcome variable, as a function of increasing number of trees per forest, over 20 iterations at each level. 2 datasets with interaction terms between features in the outcome-generating generating mechanism are included in the training set, and 2 are included in the testing set. Across all parameter levels, Weighting Trees outperforms all other approaches, and the relationship between approaches is conserved as the number of trees per ensemble increases.

Figure S2 displays Average RMSE's of the ensembling approaches as a function of increasing number of trees per forest in each ensemble. The data was generated in the same manner as described in the Simulating Datasets section of the main text, with 10 datasets used for training and 5 for validation at every iteration. Therefore, if $n$ equals the number of trees per forest, the total number of trees per ensemble is $n \times 10$. We tested ensembles with total number of trees ranging from 100 to

7500, in order to determine how the relationship between the approaches is affected by increasing ensemble size. Due to the great computational time and resources required for training the larger forests, we conducted 20 iterations (instead of the typical 100) at each forest size, resulting in larger confidence bands. We find that overall, the relationship between ensembling approaches mirrors that from Figure 2 of the main text, with Weighting Forests outperforming the others throughout. Furthermore, we see that these patterns remain constant as ensemble size increases.

There is a steep improvement in performance for the smaller ensembles as the number of trees per forest increases, but the performance of all approaches plateaus for larger ensembles; for instance, there does not appear to be significant improvements from using 750 trees per forest over 100 trees per forest. The vast increase in computational time necessary to train larger ensembles does not seem to result in appreciable gains. These findings motivate our choice of using 10 trees per forest in our analyses.
